# Integration of localized microbiome, metabolome, and clinical datasets predicts healing in chronic wounds among veterans

**DOI:** 10.1101/2025.09.07.674782

**Authors:** Catherine B. Anders, Hannah L. Smith, Jeremy Boyd, Michael C. Davis, Tyler M.W. Lawton, Chiachi Hwang, Margaret M. Doucette, Mary Cloud B. Ammons

## Abstract

The chronic wound microenvironment consists of a complex milieu of host cells, microbial species, and metabolites. While much is known about wound microbiomes, our knowledge of metabolic landscapes influencing wound healing is limited. Furthermore, integrating complex datasets into predictive models of wound healing is almost non-existent. Microbial rRNA and total metabolites were extracted from 45 diabetic foot ulcers (DFU) debridement samples from 13 patients, with 25 from non-healing wounds and 20 from healing wounds that remained closed for over 30 days. 16S rRNA sequencing and global metabolomics were performed and clinical metadata collected. Healing outcome was modeled as a function of three blocks of features (N = 21 clinical, 634 microbiome, and 865 metabolome) using DIABLO (**D**ata **I**ntegration **A**nalysis for **B**iomarker Discovery using **L**atent C**o**mponents). The final model selected 176 features (N = 15 clinical, 8 microbiome, and 153 metabolome) and the correct clinical outcome was predicted with an accuracy of nearly 94%. These results indicate that integrating multi-omics data with clinical metadata can predict clinical wound healing with low error rates. Furthermore, the biomarkers selected within the model offer novel insights into wound microenvironment composition which may reveal innovative therapeutic approaches and improve treatment efficacy in difficult-to-heal wounds.

## INTRODUCTION

The number of people over the age of 18 diagnosed with type 2 diabetes mellitus (T2DM) quadrupled from 5.5 million to 21.9 million from 1980 to 2014, representing 9.1% of the adult population (1). These numbers are projected to increase to 39.7 million (13.9 %) in 2030, and 60.6 million (17.9%) in 2060 (1). Within this affected population, it is estimated that up to 44% will develop a diabetic foot ulcer (DFU) infection (2), which leads to amputation in approximately 25% of this population due to failure to heal (3). Independent risk factors associated with the development of DFU infections and amputation are variable, and include peripheral neuropathy, peripheral arterial disease (PAD), periwound edema, wound area, wound depth, elevated C-reactive protein, polymicrobial infections, extended spectrum beta-lactamase-producing Gram-negative bacterium, hospital admission for DFUs, and vancomycin treatment (3–6). Given that the 5-year mortality rates following lower extremity amputations are approximately 50% (7), a clearer understanding of the underlying factors impacting healing outcomes of DFUs is needed.

Bacterial bioburden, biofilm formation, and recurrent infections have been identified as contributing factors to the development of chronic wounds—generally defined as wounds that persist 4-6 weeks post wounding and do not respond to standard treatment regimens (8). It is estimated that 50% of chronic DFUs will be infected upon examination, but fail to demonstrate clinical signs of infection due to PAD or patient immune system dysregulation (9). While the traditional wound culture methods employed by most diagnostic labs can identify the presence of pathogenic bacteria, this information may offer little diagnostic value as all open wounds will colonize bacteria, and culture-based methods are limited to bacteria that will grow well in culture and often underestimate the true bacterial diversity compared to culture-independent methods (10, 11). Previous studies focused on culture-based methods have identified *Staphylococcus aureus*, *Enterococcus faecalis*, *Enterobacter cloacae*, *Pseudomonas aeruginosa*, *Acinetobacter baumanni*, *Escherichia coli*, *Corynebacterium* species (spp.), *Klebsiella* spp., and *Streptococcus* spp. (4, 12, 13) as pathogens that contribute to infections within the wound microenvironment. While culture-based methods can identify pathogenic bacteria that will grow well in culture, these methods are limited in their ability to detect less abundant, non-pathogenic bacteria and anaerobic species which could significantly influence healing (11). Culture-independent techniques, such as 16S ribosomal RNA (rRNA) and next generation sequencing (NGS), allow for a more comprehensive assessment of microbiota diversity that enable the use of complex statistical analyses capable of identifying predictors that influence healing progression, including the assessment of therapeutic treatment, subtle changes in microbial diversity over time, the presence of biofilm associated pathogens, and bacteria associated with patient metabolic status (12, 14–18).

Within chronic wounds, colonizing bacteria exist in viable, dormant, and non-viable physiological states with pathogenic bacterial species demonstrating high metabolic activity while metabolically dormant species contribute to chronic wound persistence (19). It has been hypothesized that concentration gradients of key metabolites established during bacterial colonization and biofilm formation drive the activation of these variable metabolic states (20); however, few studies have sought to characterize the metabolome of chronic wounds. Ammons, et al. (19) examined the microbiome and metabolome of chronic pressure ulcers and determined that distinct shifts in the metabolome of these wounds corresponded to changes in the microbiota present.

Likewise, a temporal healing study of acute wounds revealed that decreases in *Staphylococcus aureus* corresponded to increases in *Propionibacterium* and collagen levels within the wounds (21). These studies indicate that it is particularly important to identify how the metabolome of the chronic wound microenvironment modulates the composition of the colonizing bacteria, and how those shifts impact patient healing.

While the contribution of colonizing microbiota and localized metabolic environment are known to act synergistically with patient predisposition to drive clinical outcome in wound healing, to our knowledge there are no studies that combine such data into a single predictive model of clinical outcome. In the present work, we utilized a leading method for handling the integration of high-dimensional data: DIABLO (**D**ata **I**ntegration **A**nalysis for **B**iomarker Discovery using **L**atent C**o**mponents) (22). DIABLO is a multi-dataset relative of sparse partial least squares discriminant analysis (sPLS-DA) that learns a multi-omics signature capable of predicting whether chronic wounds will heal. The model was employed in two ways. First, the model itself — the features selected from each dataset and the pattern of correlations among features — was used to generate systems-level hypotheses regarding why some wounds in the study population heal while others do not. Second, selected metabolome and microbiome features were fed into additional non-integrative analyses: metabolite set enrichment analysis (MSEA), and investigation of relative bacterial abundances by outcome class.

It is worth noting how the largely integrative focus taken here differs from traditional univariate biomarker discovery. Univariate methods test individual features one at a time to see whether they are significantly associated with an outcome. In contrast, the present study’s integrative model identifies clusters of correlated features predictive of wound healing. This correlative structure reflects systems-level interactions between host health, metabolome, and microbiome, and offers a rich set of constraints on hypothesis generation.

In sum, this study addresses the need for the comprehensive characterization of both the microbiome and the metabolome of chronic wounds and correlates those findings to healing outcome and patient metadata. Our results demonstrate the usefulness of a multi-omics modelling approach, and provide a foundation for development of novel, clinically impactful wound healing diagnostics and therapeutics.

## RESULTS

### Patient Cohort Characterization (Table 1)

For this study, the patient cohort was recruited from the Boise VA Wound Clinic and selected for having wounds recalcitrant to acute healing, as defined in the Methods. This cohort is older, Caucasian, male, and has a higher rate of comorbidities compared to the general population. The average age for the cohort was between 69 – 80 years old. Overall, the donor cohort was predominantly diabetic (33% of Healers, 71% of Non-Healers) and prediabetic (50% of Healers, 14% of Non-Healers). Non-diabetic patients accounted for a minority of the healing and non-healing cohorts (16% and 14%, respectively). Nearly the entire cohort was classified as overweight or obese based on Body Mass Index (BMI 25), with no statistically significant difference noted between the Healers and Non-Healers. In addition, the non-healing cohort was notably hyperglycemic with a statistically significant increase in HA1C compared to the Healers which was reflected by comparable differences in overall glucose levels between the two groups. Clinical detection of antimicrobial resistant pathogens (ARP) within colonizing wounds was found to be statistically significant between the healing and non-healing groups, with significantly higher levels of ARP found in Non-Healers. The other marker of overall health found to be statistically significant between the healing and non-healing groups was monocyte percentage. Associated comorbidities were slightly higher in the non-healing cohort .

### Integrative Model

The final, two-component DIABLO model selected 176 of the 1,055 features submitted. This was sufficient to achieve a cross-validated error rate of 6.4%, meaning that the model was able to correctly classify unseen samples with nearly 94% accuracy. A one-component version of the final model had a 10.7% error rate, which simultaneously demonstrates the increased predictive accuracy associated with the addition of a second component, and argues against adding a third component, since performance improvements would likely be marginal. Supplemental Table 1 summarizes the number of features selected into the final model by dataset and component.

One way that the model-derived multiomic signature can be represented is as a network graph. In Figure 2 for example, selected features that correlate with features in other datasets above/below ±0.45 are depicted as points. Color denotes the dataset each feature belongs to, and shape indicates higher expression in healing versus non-healing wounds. The graph representation captures the model’s two-component solution.

**Figure 1:**
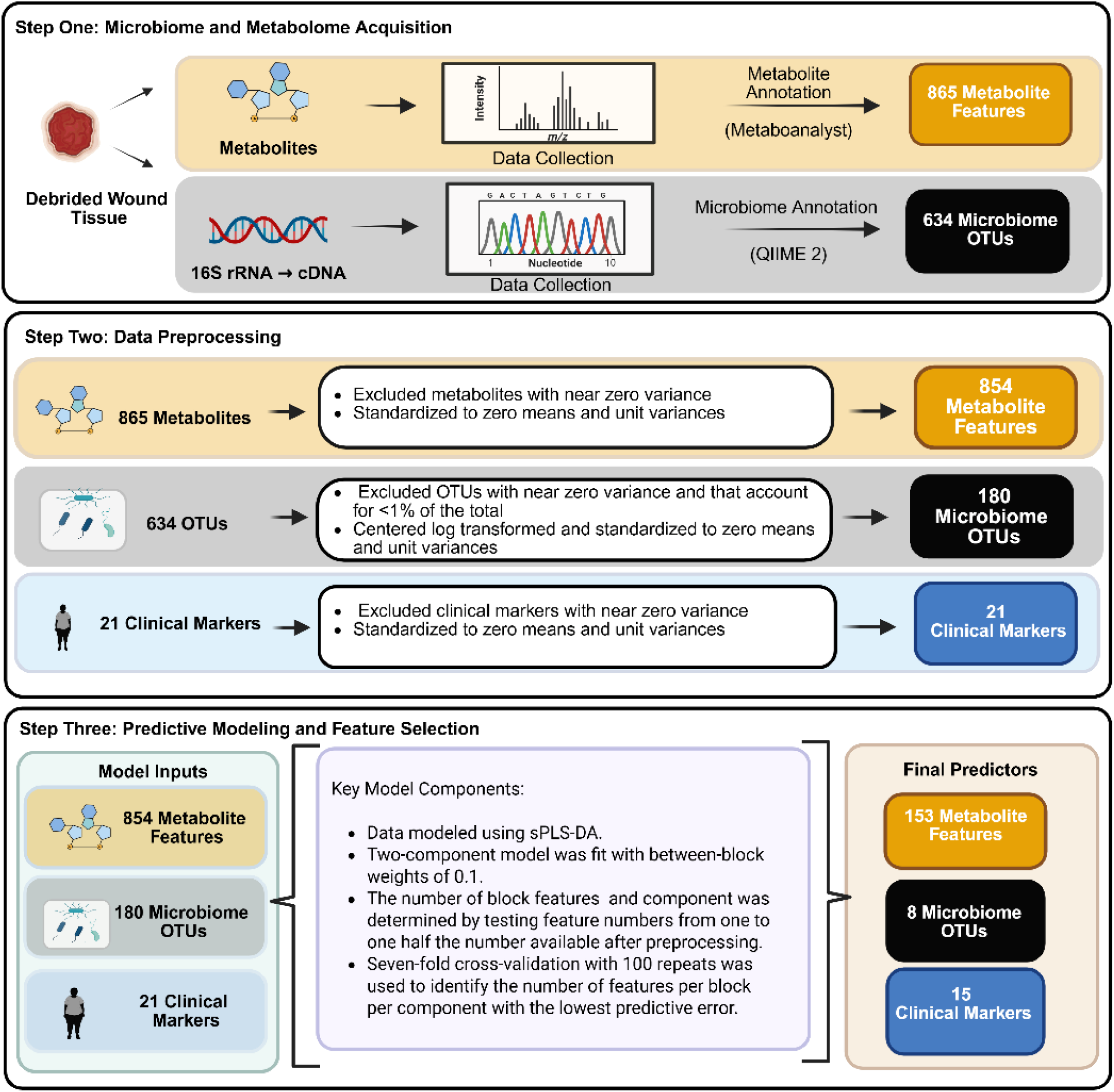
Analytical overview of the predictive model using the metabolome, microbiome, and clinical marker feature sets. First, localized chronic wound metabolite profile and colonizing microbiome OTUs were collected and annotated utilizing Metaboanalyst 5.0 and QIIME 2. Second, data preprocessing steps were applied to eliminate near zero variance and to standardize the metabolome, microbiome, and clinical marker feature sets. Third, the predictive model was applied to select features that could predict healing status with the lowest error rates.

**Figure 2:**
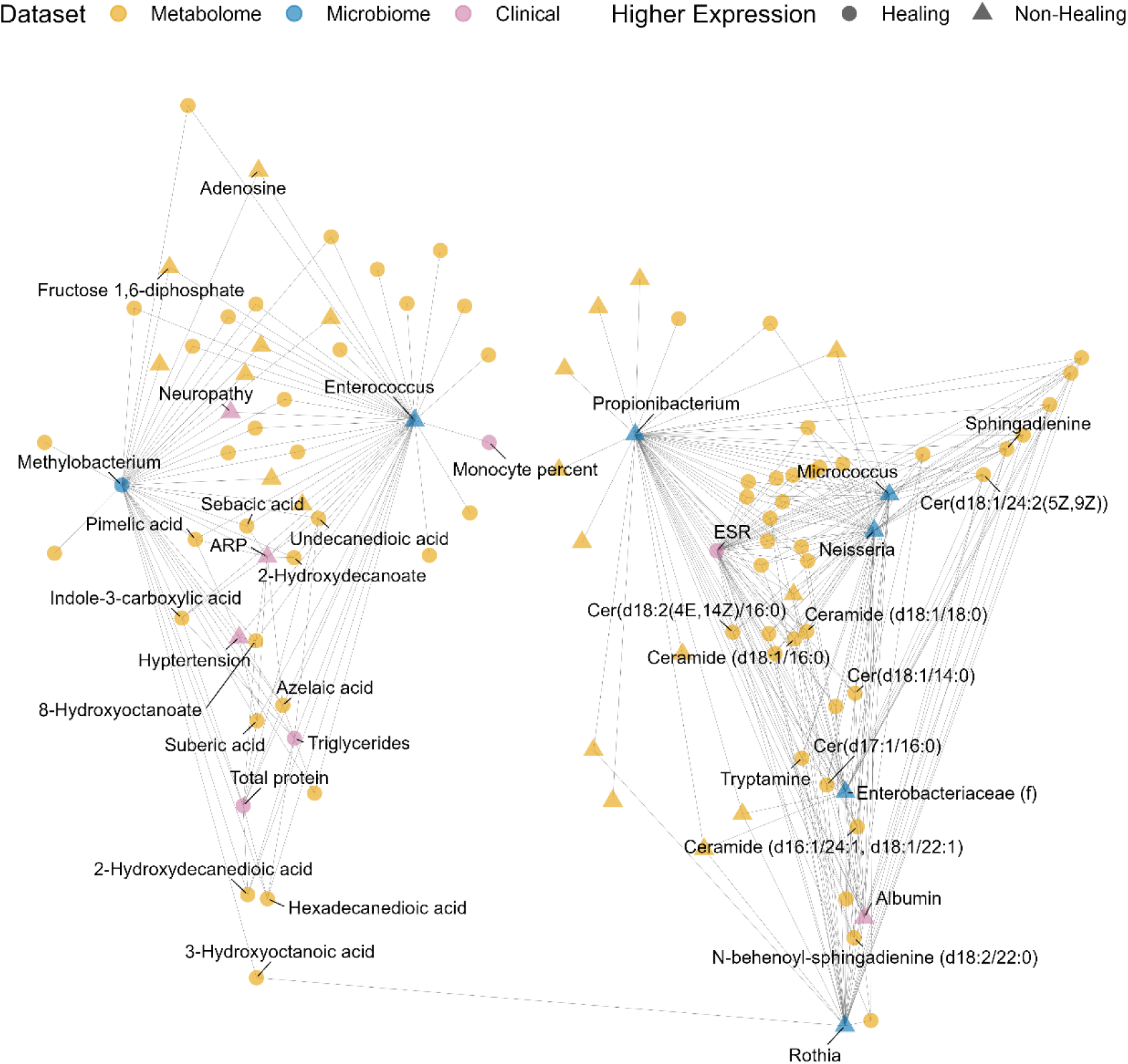
The two-part multi-omics signature discovered by the model can be depicted by a network graph. Component one (left cluster) and component two (right cluster) of the model can be visualized within the network graph. Each point in the graph represents either a metabolome (orange), microbiome (blue), or clinical (purple) feature. Features with higher expression in healing wounds are plotted as circles; those with higher expression in non-healing wounds are plotted as triangles. Only model features with at least one correlation to another feature above/below ±0.45 are shown.

Features that are more heavily loaded on component one—like *Enterococcus*, *Methylobacterium*, and associated metabolites and clinical features—are shown in the group to the left. Whereas features that are more heavily loaded on component two— such as *Propionibacterium*, *Neisseria*, *Micrococcus*, and assorted ceramides and sphingadienines—appear on the right.

Additional structure in the model’s solution can be viewed in Figure 3, which plots all 176 selected featues as points in two-dimensional space according to their correlation with components one (*x*-axis) and two (*y*-axis). The location of each feature in the figure encodes three critical types of information. First, features further from the origin are more important for predicting the outcome: those closer to the outer dashed circle (which represents correlations of ±1) are more heavily weighted in the model than those near the inner dashed circle (which indicates correlations of ±0.5). Second, the angle created by vectors drawn from any two features through the origin gives the sign of the correlation between features: acute, obtuse, and right angles correspond to positive, negative, and null correlations, respectively. Third, the length of the vectors signifies the magnitude of the correlation. For example, two points close to each other and to the outer dashed circle are more strongly positively correlated than two points whose vectors form the same angle but are near the inner circle.

**Figure 3:**
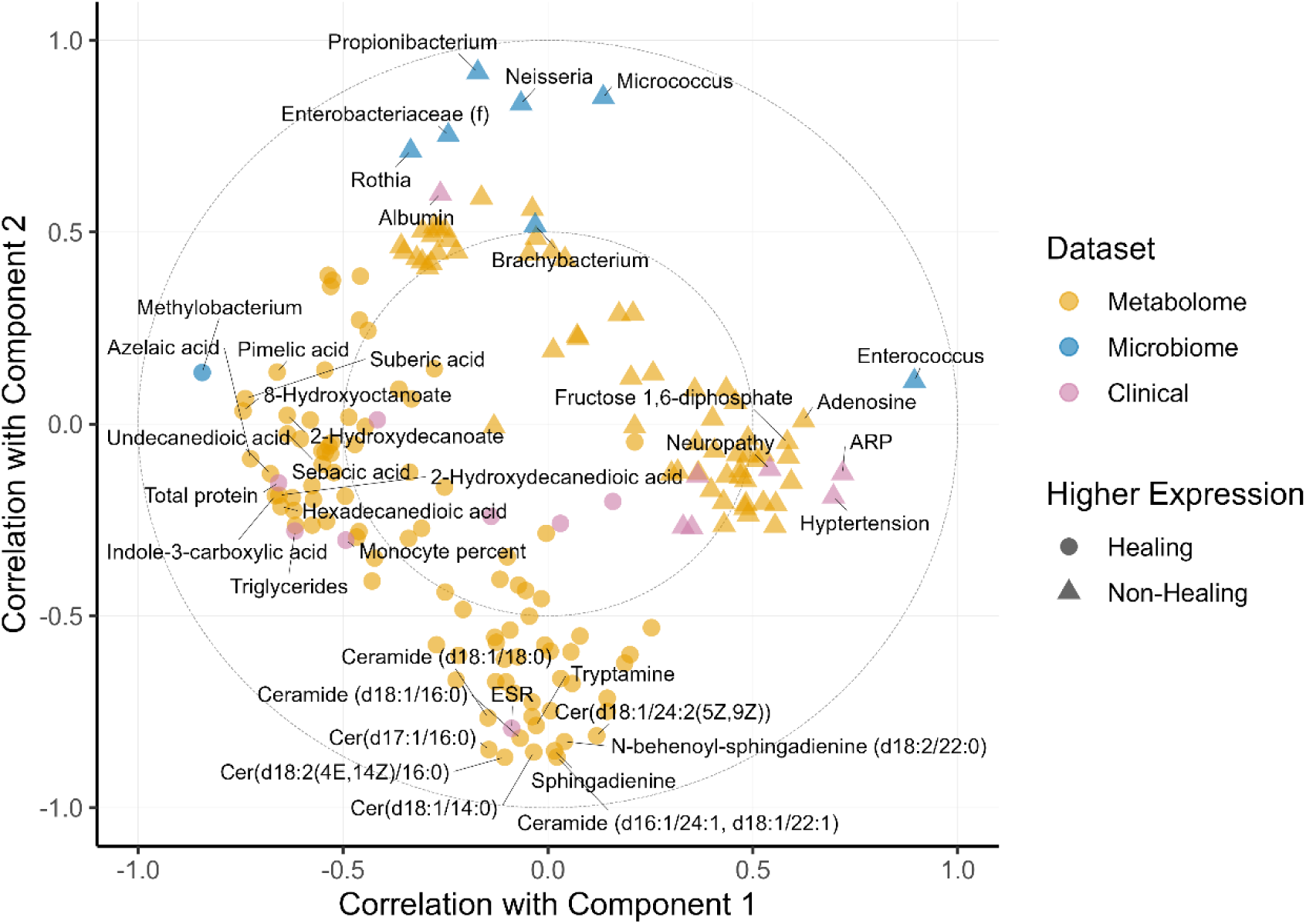
Distinct feature clusters can be identified from the features selected into the final DIABLO model. The Circle plot illustrates both the predictive two-part multi-omics signature discovered by the model and feature clusters within each model component. All 176 features selected into the model are located in 2D space according to their correlations with model components one (x-axis) and two (y-axis). The large dashed circles are guides representing correlations of ±0.5 (inner circle) and ±1 (outer circle). Colors indicate whether selected features belong to the metabolome (orange), microbiome (blue), or patient marker (purple) data sets. Shapes indicate higher expression in healing wounds (circles), or non-healing wound (triangles). Four clusters of related features can be identified: Cluster 1 (left) is centered around *Methylobacterium*; Cluster 2 (right) is centered around *Enterococcus*; Cluster 3 (bottom) centers around a series of metabolite features; and Cluster 4 (top) centers around a group of microbial features.

Whereas Figure 2 divides the features according to component, Figure 3 allows further categorization into four distinct clusters — Clusters 1 and 2 (C1 and C2) are composed of features with the most negative and positive correlations to component one, while Clusters 3 and 4 (C3 and C4) have the most negative and positive correlations to component two. Clusters are also differentially associated with outcomes. For instance, C1 is composed of features that have higher expression in healing wounds: *Methylobacterium*, the metabolites 8-hydroxyoctanoate, suberic acid, and azelaic acid, and clinical measurements for total protein and triglycerides. C2 includes features that have higher expression in non-healing wounds: *Enterococcus*, compounds like adenosine and fructose 1,6-diphosphate, and clinical markers indicating ARP, hypertension, and neuropathy. C3 contains ceramides, sphingadienines, tryptamine, and ESR (the erythrocyte sedimentation rate), all of which have higher expression in healing wounds. The features in C4 have higher expression in non-healing wounds— e.g., the bacteria *Propionibacterium*, *Neisseria*, and *Micrococcus*—and the clinical measure for albumin.

Importantly, features in the same cluster are positively correlated with one another and negatively correlated with features in the opposite cluster. These relationships are made explicit in Figures 4 and 5. Figure 4 illustrates pairwise correlations between features with the 30 largest component one loading weights. Light colored blocks indicate positive correlations within C1 and C2, whereas dark blocks indicate negative cross-cluster correlations. The same pattern is evident in Figure 5: features in C3 and C4 are positively correlated with features in the same cluster but negatively correlated with features in the opposite cluster.

**Figure 4:**
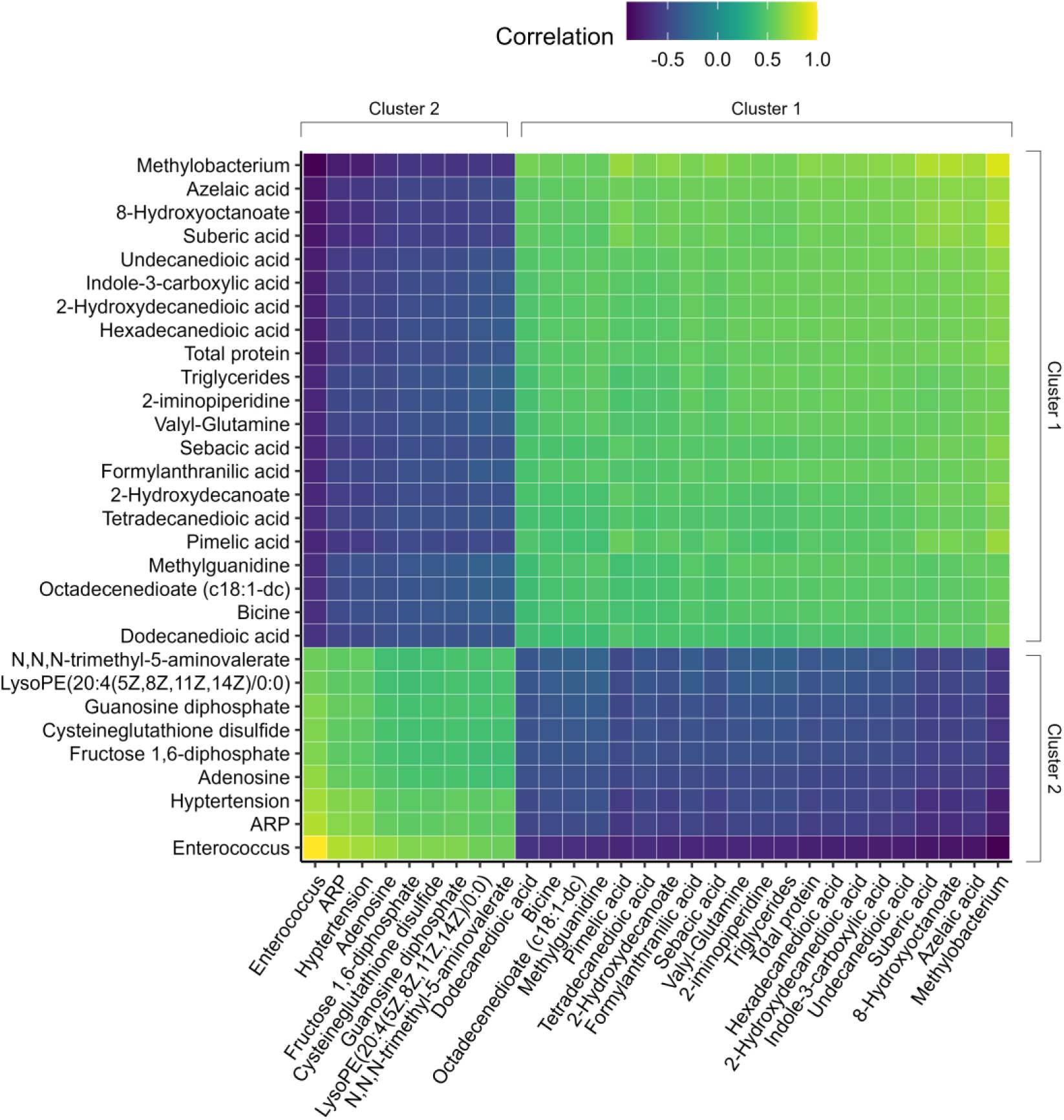
Heatmap demonstrates that features are positively correlated within clusters and that cluster 1 is negatively correlated to cluster 2. Heatmap depicting pairwise correlations between features with the 30 largest component one loading weights. Correlations are the dot product of the vectors created by joining two features through the origin in Figure 3. All correlations are normalized to the value of the feature pair with the longest vectors, which in this case is the correlation between *Enterococcus* and itself.

**Figure 5:**
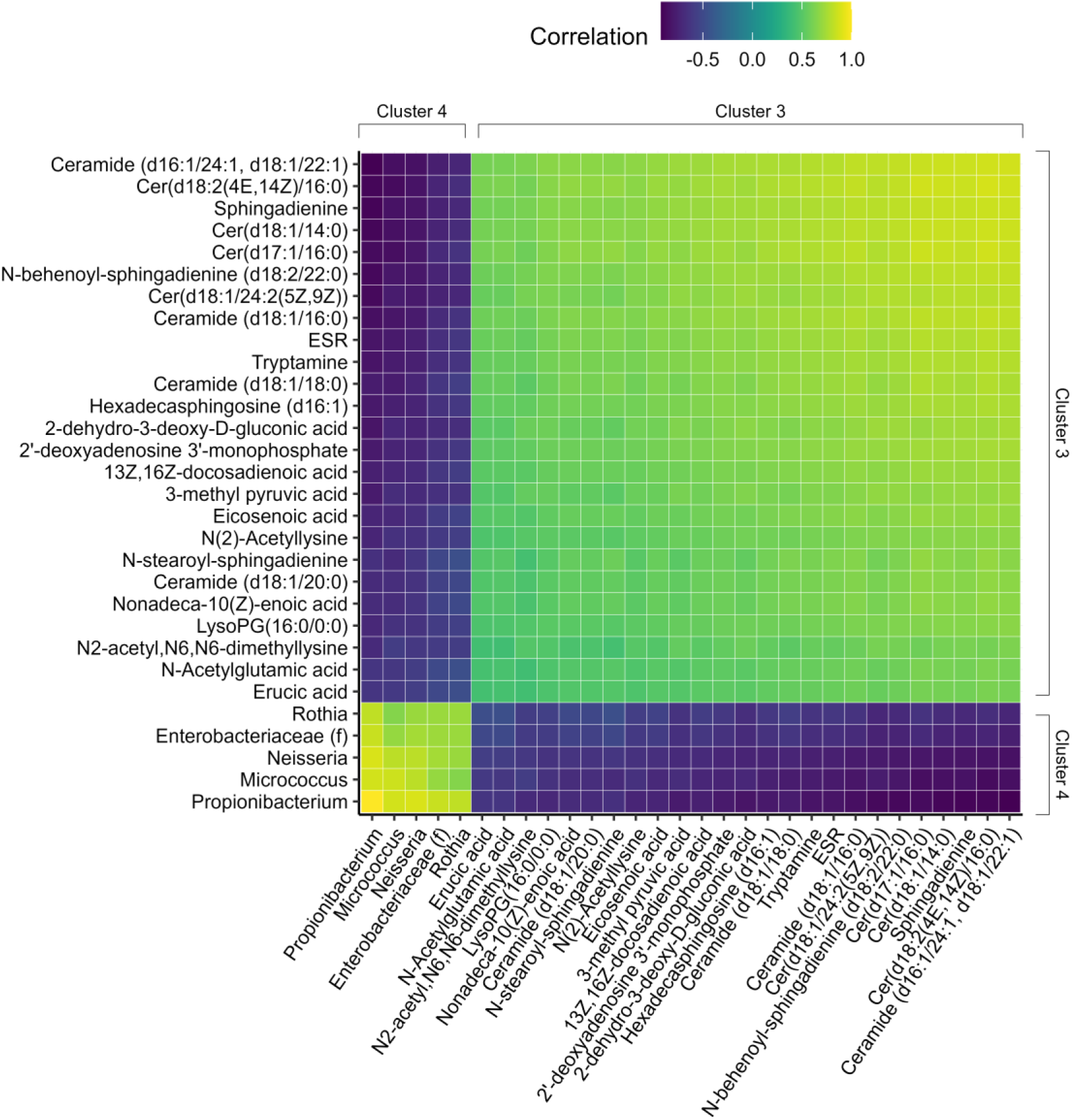
Heatmap demonstrates that features are positively correlated within clusters and that cluster 3 is negatively correlated to cluster 4. Heatmap depicting pairwise correlations between features with the 30 largest component two loading weights. Correlations are the dot product of the vectors created by joining two features through the origin in Figure 3. All correlations are normalized to the value of the feature pair with the longest vectors, which in this case is the correlation between *Propionibacterium* and itself.

### Characterization of selected metabolites

From the metabolomics data set, 153 metabolites were selected by the DIABLO model with 95 associated with Healers and 58 correlated to Non-Healers. Metabolite subclass analysis was utilized to identify key metabolic constituents of healing and non-healing wounds. For the purposes of this analysis, chemicals and drug metabolites that were associated with patient medications were excluded from the subclass analysis leaving 91 healing and 56 non-healing metabolites represented in Figure 6. For Healers (Figure 6A), fatty acids represented the most abundant metabolite subclass at 36.4% (32/91) while amino acid and peptides were more abundant in Non-Healers (Figure 6B) at 21.4% (12/56). The highest contributing features to the model besides fatty acids within healing wounds were amino acids and peptides and ceramides at 17.0% (15/91) and 13.6% (12/91), respectively (Figure 6A). Within non-healing wounds, fatty acids and purines were the next two most abundant subclasses at 21.6% (11/56) and 11.8% (6/56), respectively (Figure 6B).

**Figure 6:**
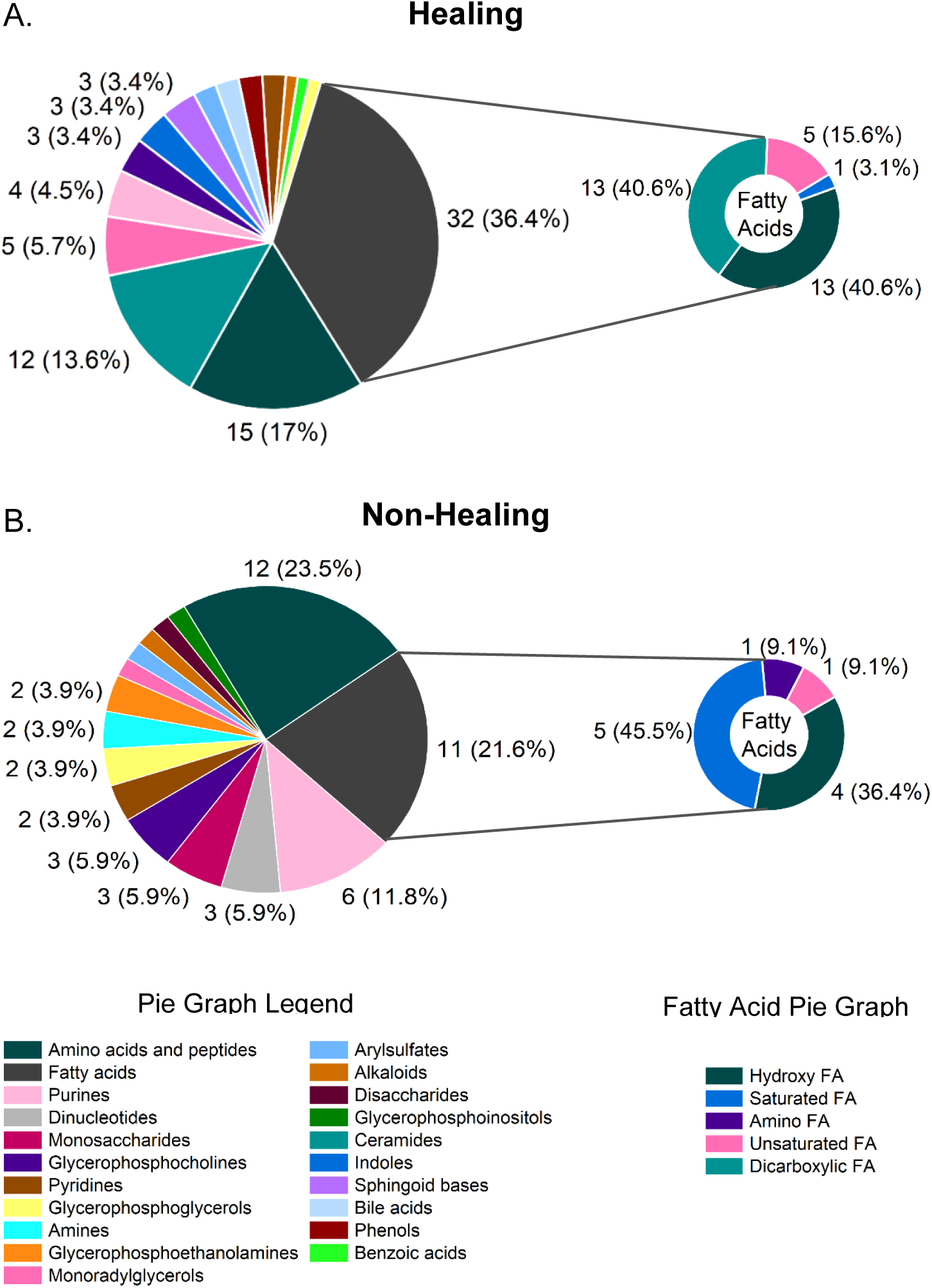
Healing and non-healing wounds contain disparate metabolic environments. Classification of the 153 metabolome features selected by the DIABLO model were classified by metabolic subclass for (A) Non-healing wounds and (B) Healing wounds. Pie chart annotations indicate the number (percent) of selected features making up each category. The top 15 subclasses are shown for each healing outcome. Metabolite subclasses comprising <3% of the total are depicted but not labeled on the pie chart.

Within the broader metabolite set, both healing and non-healing wounds contained high percentages of fatty acids; however, when fatty acids are considered as a standalone subclass, 74.4% (32/43) of the fatty acids are more abundant in healing wounds compared to only 25.6% (11/43) for non-healing wounds. Additionally, there were notable differences between the patient groups in the fatty acid subclass composition (small pie graphs, Figure 6). For example, all dicarboxylic fatty acids — 40.6% of the healing group fatty acids — selected into the model had higher expression in healing wounds, while no dicarboxylic fatty acids were present in non-healing wounds. Within the non-healing group, 45.5% of fatty acids were saturated, but only 3.1% saturated fatty acids were detected within healing wounds. No amino fatty acids were selected for the healing category, while 9.1% of the fatty acids in the non-healing wounds were amino fatty acids. Differences within the amino acid subclass were equally disparate between the two healing outcomes. In Healers, 3/15 or 20% of the amino acid metabolites were dipeptides while the remaining metabolites represented eight different amino acid metabolic pathways including alanine, tyrosine, glycine, serine, threonine, histidine, lysine, glutamate, arginine, proline, and branch-chain amino acid pathways.

Within Non-Healers, histidine and lysine pathways metabolites comprised 16.7% (2/12) and 33.3% (4/12), respectively, of the pathways represented. Finally, metabolic classes represented solely in Healers consisted of ceramides (13.6%), indoles (3.3%), sphingoid bases (3.3%), bile acids (2.2%), phenols (2.2%), and benzoic acids (1.1%). Within non-healing samples solely representative subclasses included dinucleotides (6.7%), monosaccharides (6.7%), amines (3.3%), pyrimidines (3.3%), disaccharides (1.7%), and short-chain acids (1.7%).

To identify the impacted pathways, the metabolic features selected by the model for each clinical outcome were then analyzed for metabolic pathway impact by metabolite set enrichment analysis (MSEA). In total, 63 metabolic pathways were identified for Healers and 39 were selected for Non-Healers. Categorically, most of the metabolic pathways in both Healers and Non-Healers were associated with amino acid and energy metabolism. In Healers, amino acid and energetic pathways accounted for 31.7% and 28.6%, respectively, of the represented pathways compared to 46.2% and 35.9% for amino acid and energy metabolism, respectively, within non-healing samples.

The most significant difference between the two sample sets was the prevalence of fatty acid pathways with 19.1% noted for Healers compared to 5.1% for Non-Healers. The top 20 impacted pathways for each patient cohort are illustrated in Figure 7 with a complete list included in Supplemental Data Figure 7 tab. For the healing outcome, significantly impacted pathways included sphingolipid metabolism, beta oxidation of long chain fatty acids, and caffeine metabolism. Of these sphingolipid and caffeine metabolism pathways were observed exclusively within Healers. For the non-healer outcome, significantly impacted pathways included beta oxidation of long chain fatty acids, fatty acid biosynthesis, histidine metabolism, citric acid cycle, gluconeogenesis, purine metabolism, and the Warburg effect. Among fatty acid pathways, sphingolipid, phosphatidylethanolamine, glycerolipid, Phosphatidylcholine, and phosphatidylinositol phosphate metabolic pathways were identified exclusively with Healers. Other exclusively enriched pathways within healing wounds included biotin, lactose degradation, phenylacetate, thiamine, trehalose degradation, and tryptophan metabolic pathways. Only four pathways were exclusively enriched in non-healing wounds – methylhistidine, beta oxidation of saturated fatty acids, homocysteine degradation, and pyruvaldehyde degradation pathways.

**Figure 7:**
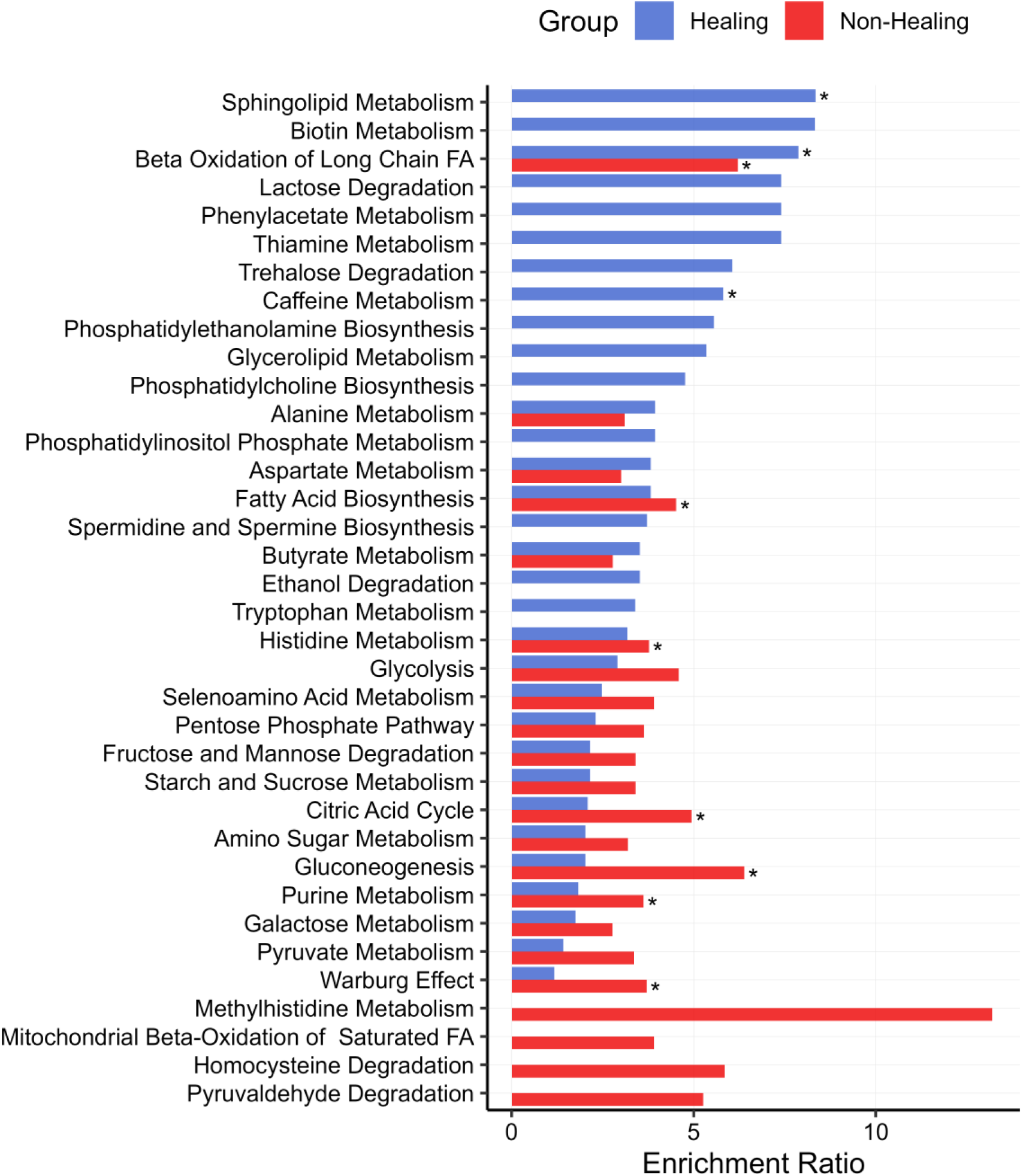
Metabolite set enrichment analysis (MSEA) of metabolite features selected by the DIABLO model demonstrate differential impacted metabolic pathways in healing and non-healing wounds. Metabolic pathways impacted by feature selection are shown for healing (Blue) and non-healing (Red) wounds. The top 20 enriched pathways in each group are shown. Asterisks designate significantly enriched pathways.

### Characterization of selected microbiome features

As with metabolite features, the DIABLO model identified predictive microbiome features associated with clinical outcomes. Of the 634 taxa identified in the full dataset, 80 microbial features were introduced into the model after data pre-processing. These taxa, identified down to the genus level, when possible, are shown in a heat tree diagram (Figure 8), with relative abundance indicated by the color range. Branches and nodes colored toward blue are more abundant in Healers, while those colored toward red are more abundant in the non-healing samples. In this phylogenetic heat tree, the eight microbial features selected by the model are indicated by bold text. This heat tree highlights both the major players in the integrated model, as well as commonly cited microbes that are often regarded as major players contributing to non-healing but were not selected by the model as predictive of wound response to standard of care. Note that the two major microbial contributors to model separation along component one, *Methylobacterium* and *Enterococcus*, are clearly shown to have opposite abundance in healing versus non-healing wounds, respectively. Other microbial features selected by the model and found in component two include *Neisseria, Enterobacteriaceae*, *Brachybacterium, Propionibacterium*, and *Micrococcus* (Figure 8).

**Figure 8:**
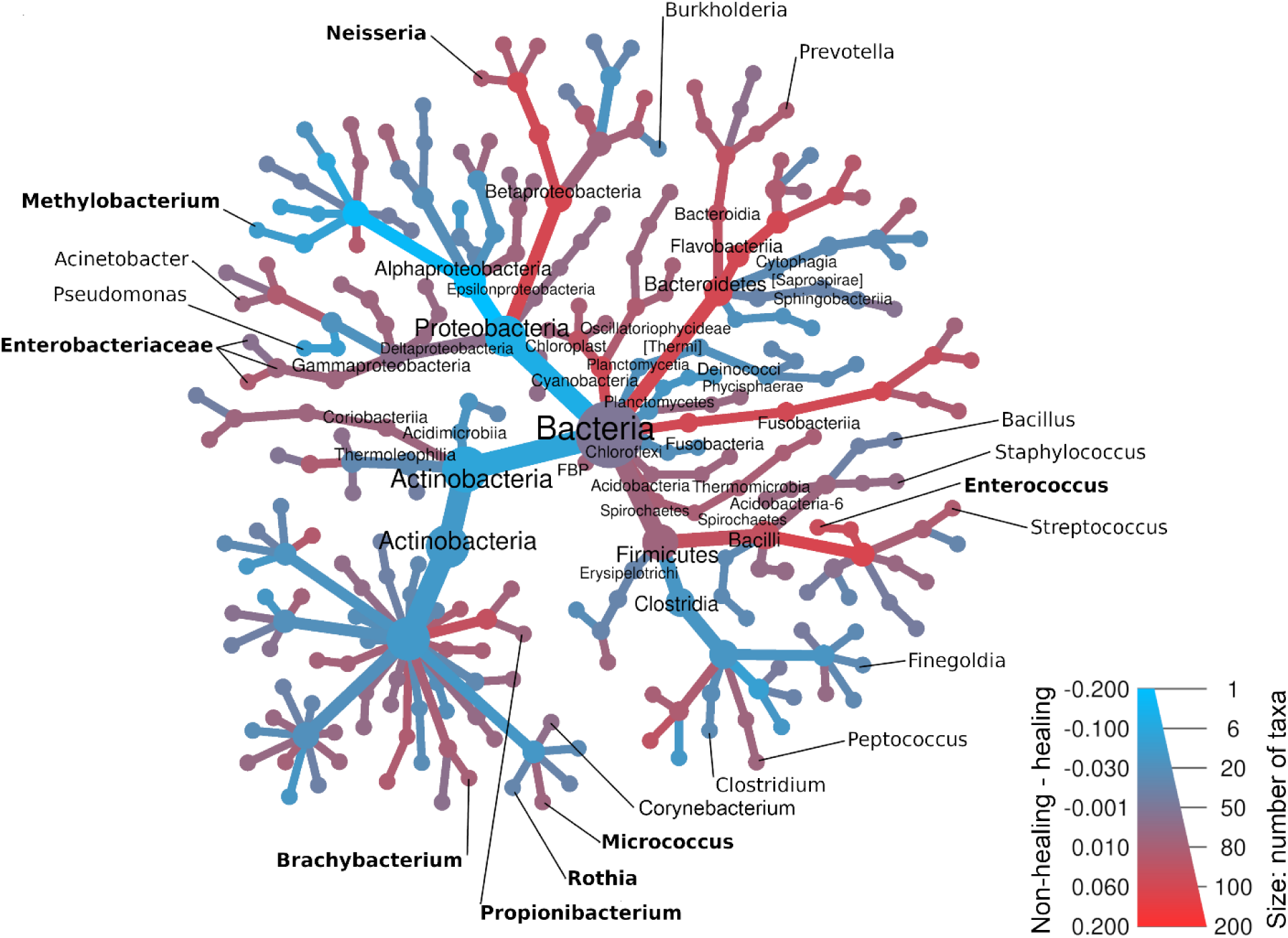
Phylogenetic heat tree of microbial features highlights microbiome differences between healing and non-healing wounds. Heat tree showing relative abundance and phylogenetic relationships among 180 taxa input into the model, between samples from healing and non-healing wounds. Nodes size indicates the number of taxa, while color denotes taxa that are more abundant in healing (blue) versus non-healing (red) wounds. Microbial features selected by the integrated model are indicated in bold.

Based on relative abundance (Supplemental Figure 1), at the phylum level, the three main phyla present in both healing and non-healing wounds were Firmicutes, Actinobacteria, and Proteobacteria, which is consistent with previous wound healing studies. Bacteroidetes was the fourth most common phylum observed, having ≈1.5% abundance across all samples. Phylum Firmicutes had greater abundance in non-healing samples at 63% compared to healing samples at 49%. As expected, *Enterococcus* (C2) is a member of the Firmicutes phylum along with other common wound associated pathogens including *Staphylococcus*, *Streptococcus, Clostridium, and Propionibacterium* (C4) *genera.* Conversely, healing wounds contained more Proteobacteria than non-healing wounds with 23% and 9% relative abundance, respectively. *Methylobacterium* (C1), *Neisseria* (C4), and *Enterobacteriaceae* (C4) all belong in these phyla with the remaining C4 microbiome features belonging to the phylum Actinobacteria.

## DISCUSSION

The primary objective of the present study was to inform understanding of wound healing in the study population by building a model that integrates across metabolome, microbiome, and clinical datasets. Our resulting DIABLO model has excellent predictive performance, correctly classifying 93.6% of held-out samples. At the heart of the model is information about how the set of 176 selected features correlate not only with the outcome, but with one another. This knowledge is the key to integration and allows essential constraint of the space of hypotheses that can reasonably explain — at the systems level — why some chronic wounds heal while others do not.

The model was oriented across two separate components. component one, arrayed along the x-axis of the circle plot (Figure 3) contained two distinct clusters, C1 (Figure 3 right) associated primarily with Healers and C2 (Figure 3 left) corresponding with Non-Healers and highlighted the inverse relationship of *Methylobacterium* and *Enterococcus*, respectively. Arrayed along the y-axis, component two encompassed the healing associated C3 and the non-healing centered C4 containing the ceramide metabolites (Figure 3, bottom) contrasted with the remaining microbiome features of the model in C4 (Figure 3, top).

*Methylobacterium* (C1) is an interesting finding of this study, worthy of follow-up in future studies. *Methylobacterium* is commonly encountered in human gut microbiome studies (23), and is a common soil and rhizosphere (plant root associated) bacterium, and has been found in relatively large quantities in other vertebrate gut microbiomes (24). Despite its general classification as an environmental commensal bacterium (25), the presence of *Methylobacterium* in healing wounds may suggest a benign microbiome (26, 27). Although *Methylobacterium* has been suspected of being a contaminant in some microbiome studies (28), its non-random distribution in our samples, showing up in appreciable amounts almost exclusively in Healers, points to it being an actual constituent of the microbiome of these wounds.

Metabolically, C1 contained an abundance of medium chain and long chain dicarboxylic acids (DAs; e.g., pimelic, suberic, azelaic, sebacic, dodecanedioic, undecanedioic, tetradecanedioic, hexadecanedioic, plus 8-hydroxyoctanoate) and methylxanthine and alkaloid metabolites (caffeine, theophylline, trigonelline, and vanillin) . DAs are a biproduct of active ω-oxidation followed by peroxisomal β-oxidation metabolic pathways, present within the MSEA for the healing group, and facilitate lipid cleanup and fatty acid oxidation when cells are under lipid metabolic stress (29). The resulting DAs, especially azelaic acid, exert antimicrobial and anti-inflammatory actions in skin (29–31). The co-occurrence of the total protein and monocyte percent clinical markers within this healing cluster suggests a better nourished, immune-competent state where macrophages can orchestrate the transition from inflammation to repair, and higher triglycerides likely reflect available energy substrates needed for remodeling rather than the maladaptive dyslipidemia seen in non-healers (32–35). A second, highly distinctive element in C1 is the methylxanthine and alkaloid signature which is consistent with caffeine metabolic pathways. These features may contribute to a healing environment via systemic benefits as habitual caffeine intake has been linked to better insulin sensitivity and lower inflammation (36–38), and through local signaling pathways which temper inflammation (39–41).

*Enterococcus* species, the dominant feature in the non-healing associated C2, are known prolific biofilm formers in chronic wounds that contribute to slow healing, resist antibiotic treatment, modulate host immunity, and sustain a persistent low inflammatory state (42–44). Mechanistically, this is supported by metabolomic and clinical features that indicate a wound environment driven by hyperglycemic-fueled metabolic stress and non-healing. The clinical marker profile, characterized by elevated ESR, high HbA1c, neuropathy, hypertension, and antibiotic-resistant organisms, reflects the classic high-risk DFU phenotype (45–47). This high-risk DFU phenotype was also evidenced by the presence of D-ribose (48), 2-aminoadipic acid (49), and 5-(galactosylhydroxy)-L-lysine (50) all of which are indicative of diabetic risk. The metabolites 2-methylbutyroylcarnitine and N,N,N-trimethyl-5-aminovalerate, which originate from microbial trimethyllysine, point towards the inhibition of carnitine biosynthesis and incomplete fatty acid oxidation (51–54). The presence of cysteine-glutathione disulfide flags oxidative stress (55). The highest enriched metabolic pathway for non-healing patients, methylhistidine metabolism, was represented by metabolite features 3-methylhistidine and N-acetyl-3-methylhistidine and is consistent with muscle/ECM proteolysis which further supports a non-healing phenotype (56, 57). Finally, the presence of chronically elevated histamine, associated with an enriched histidine metabolism pathway, contributes to vasodilation, increased vascular permeability, and localized inflammation which might indicate the presence of ineffective inflammation and persistent infection (58, 59).

Within component two, a group of metabolites, including ceramides, ceramide precursors, and sphingoid bases, represents 31% of the selected features for C3 and is consistent with the most enriched pathway in healing wounds — sphingolipid metabolism. Together, these metabolites and associated metabolic pathways are known to drive sphingolipid and ceramide metabolism that supply critical components needed for keratinocyte migration and differentiation during the proliferative and remodeling phases of wound healing (60–62). Additionally, tryptamine, a tryptophan metabolism byproduct of both microbial and host metabolism, has been shown to activate signaling receptors in keratinocytes and reduce inflammation in skin disorders to facilitate healing (63, 64). The co-occurrence of ESR, elevated in healers, is noteworthy as a systemic inflammation marker within this cluster. This trend, which has been reported in other research (65), suggests that patients with healing wounds have a higher systemic inflammatory response which is beneficial within the healing process.

Alternatively, the lower ESR levels in Non-Healers might suggest an ineffective systemic response that results in a smoldering localized inflammation that never fully engages the healing process (66).

In contrast, C4 epitomized a dysbiotic wound ecology governed by biofilm forming facultative skin colonizers including *Propionibacterium, Micrococcus, Neisseria,* and *Enterobacteriaceae*, which have the capacity to become pathogenic depending on the species present and host immunity (67–70). Fifty five percent of the C4 metabolites are medium chain fatty acids (MCFAs), including heptanoic, pelargonic, 3-Hydroxyhexanoic, caprylic, and caproic acid. The MCFAs heptanoic, 3-Hydroxyhexanoic, and pelargonic acid along with allantoic acid are associated with microbial metabolism and have been identified as potential biomarkers for impaired glucose regulation and diabetes (71–73). Additionally, many of these MCFAs have been associated with systemic or localized inflammation (74–77). Albumin, a robust marker of nutrition, normally correlates with higher total protein levels as seen in our healing patients (78); however, in this data, albumin has slightly higher levels in Non-Healers. Low albumin has been associated with delayed wound closure following surgical wound debridement or amputation (79). While the albumin levels for Non-Healers are within the normal reference range clinically, it is probable that Non-Healers remain nutritionally comprised given their low total protein levels.

Viewed through the lens of chronic wounds in T2DM, these data resolved into two largely orthogonal components. Along component one, C1 at one end defined a metabolically competent wound containing benign microbial colonizers within a nutritionally resourced and immunologically capable host. Conversely, C2 embodied the classic high-risk DFU patient with non-healing wounds infected with pathogenic biofilms and a metabolic signature of a wound stuck in breakdown mode. Translationally, the model suggests that moving a non-healing patient from C2 to C1 would involve tight glycemic control, nutritional optimization and support, and aggressive infection/biofilm management. Within component two, C3 represented a reparative wound dominated by barrier lipid remodeling and effective systemic inflammation balanced by anti-inflammatory signatures. In contrast, C4 alluded to the presence of polymicrobial biofilms and microbial metabolites that contribute to sustained inflammation. Therefore, to move a patient in C4 towards the reparative C3 might involve treatment to decrease polymicrobial infections, intervention to increase barrier lipid remodeling, and nutritional support. Importantly, the two independent, orthogonal components intimate that a patient can be well controlled metabolically (towards C1) yet can still stall locally (towards C4). Furthermore, the mechanistic story that emerges is clinically intuitive, suggesting that healing happens when host metabolism, nutrition, and immunity align within a locally pro-repair lipid and commensal microbiome environment. Conversely, chronic wounds persist when pathogenic biofilms and microbial-dominated wound biochemistry overpowers a nutritionally depleted, hyperglycemic host.

The primary objective of this research was to build an integrative model that combines metabolome, microbiome, and clinical datasets to deepen our understanding of wound healing in a specific study population. The resulting DIABLO model achieved an impressive predictive performance, correctly classifying 93.6% of held-out samples.

Central to the model is the interplay among 176 selected features, which not only correlate with the healing outcomes but also with each other, allowing us to constrain and refine the hypotheses that could explain healing mechanisms at a systems level. This study’s strengths lie in its robust integrative approach and the high predictive accuracy of the resulting model; however, there are limitations to this work, including the predominantly Caucasian and male cohort from our local Veteran population, which may not fully represent a broader demographic, and a relatively small sample size necessitating further validation. While our model provides valuable correlative insights, mechanistic studies are required to establish causation. Despite these limitations, the translational impact is significant; our findings suggest that interventions addressing glycemic control, nutritional optimization, infection management, and barrier lipid remodeling could shift non-healing wounds towards healing. Overall, this integrative approach demonstrates the potential to predict wound healing by coupling the complex wound microenvironment with patient metabolic health, paving the way for improved clinical strategies in managing chronic wounds, particularly in high-risk populations.

## METHODS

### Sex as a biological variable

The patient cohort for this study was exclusively male and reflected the local Veteran population, which is predominately Caucasian and male. We cannot speak to the relevance of these results for women.

### Ethics statement

Written informed consent was obtained from patients (ages 18-95 years old) being treated in the Wound Care Clinic at the Boise VA Medical Center (BVAMC) and enrolled in the study approved by the VA Puget Sound institutional review board (IRB, IRBNet ID:1647932). Patient demographics, data from electronic medical records, and serial wound debridement tissue discarded as part of patient standard of care were collected. After collection, samples were stored at -70°C until analysis.

### Patient Cohort

For this study, patients were recruited from the BVAMC Wound Care Clinic and were selected based on the presence of a non-acute, chronic wound, defined as being recalcitrant to healing for over 30 days. Exclusion criteria included <18 years of age, pregnancy, or nursing. Patients were followed over the course of standard of care wound treatment, and serial wound debridement samples were collected for later analysis. For the purposes of this study, a Healer is any subject who had resolved a single or multiple wounds to >80% closure without the co-occurrence of another wound and remained healed for >30 days, whereas a Non-Healer is any subject who underwent amputation due to a failure to heal or presented with multiple wounds either concurrently or consecutively over an extended period without prolonged wound resolution or healing of all wounds for a period of >30 days. Demographic and clinical data from the subject medical record was extracted to characterize the total patient cohort. We studied 13 subjects treated at the Wound Care Clinic. Subjects consented to have their serial debridement tissue profiled for microbiome colonization and local metabolite landscape, resulting in a total of 45 wound debridement tissue samples (20 samples from Healers and 25 samples from Non-Healers) characterized for microbiome and metabolome analyses (Table 1).

**Table 1:** Patient characteristics.

| Characteristic | Healing<br>N = 6<sup>1</sup> | Non-healing<br>N = 7<sup>1</sup> | p-value<sup>2</sup> |
| --- | --- | --- | --- |
| Age | 69.5 (14.4) | 78.9 (11.9) | 0.235 |
| Male | 6 (100.0%) | 7 (100.0%) |  |
| BMI | 30.8 (8.6) | 31.6 (6.0) | 0.857 |
| Diabetic | 2 (33.3%) | 5 (71.4%) | 0.286 |
| Non-diabetic | 1 (16.7%) | 1 (14.3%) | >0.999 |
| Pre-diabetic | 3 (50.0%) | 1 (14.3%) | 0.266 |
| HA1C | 5.7 (0.7) | 7.5 (1.3) | **0.032** |
| Wound Size | 3.7 (3.9) | 0.9 (1.2) | 0.198 |
| Wound Depth | 0.7 (0.8) | 0.2 (0.1) | 0.109 |
| ARP | 1.3 (1.8) | 4.6 (3.2) | **0.045** |
| Amputation | 0 (0.0%) | 3 (42.9%) | 0.192 |
| Smoker | 3 (50.0%) | 4 (66.7%) | >0.999 |
| Hypertension | 4 (66.7%) | 6 (100.0%) | 0.455 |
| Anemia | 2 (33.3%) | 2 (33.3%) | >0.999 |
| Hyperlipidemia | 4 (66.7%) | 5 (83.3%) | >0.999 |
| Neuropathy | 2 (33.3%) | 3 (50.0%) | >0.999 |
| PAD | 1 (16.7%) | 2 (33.3%) | >0.999 |
| Osteomyelitis | 1 (16.7%) | 2 (33.3%) | >0.999 |
| Glucose | 121.3 (30.3) | 187.3 (87.8) | 0.101 |
| Albumin | 3.5 (0.7) | 3.7 (0.5) | 0.682 |
| Total Protein | 7.4 (0.2) | 7.1 (0.4) | 0.099 |
| Alkaline Phosphatase | 84.9 (19.5) | 92.8 (15.0) | 0.439 |
| Monocytes (%) | 11.4 (3.0) | 7.2 (2.9) | **0.034** |
| C-Reactive Protein | 3.8 (6.0) | 1.3 (0.8) | 0.520 |
| ESR (Sed Rate) | 35.2 (18.1) | 27.7 (15.4) | 0.441 |
| Triglycerides | 175.4 (78.2) | 105.7 (37.0) | 0.086 |

<sup>1</sup>Mean (SD); n (%)
<sup>2</sup>Welch's t-test; Fisher's exact test; Wilcoxon rank-sum

### Data Set Acquisition

#### Metabolomic Profiling

##### Intracellular Metabolite Isolation

With the goal of creating a comprehensive snapshot of the chronic wound metabolome, 10-30 mg of wound debridement tissue was homogenized using four cycles of 120 seconds each at 6.0 m/s with a 300 second delay between cycles in FastPrep® lysing matrix D tubes (MP Biomedicals; Auckland, New Zealand) utilizing the FastPrep-24™ 5G Homogenizer (MP Biomedicals; Auckland, New Zealand). Samples were refrozen between each homogenization cycle with liquid nitrogen. The resulting tissue homogenate was suspended 375 µL of ice-cold 100% methanol (Honeywell; Muskegon, MI, USA) and 375 µL of ice-cold ultrapure distilled water (Invitrogen; Grand Island NY, USA) followed by the addition of 700 µL of ice-cold chloroform (Acros Organics; Thermo Fisher Scientific; Waltham, MA, USA). The tubes were homogenized again using the same settings. The homogenized samples were centrifuged at 16,000 x g for 5 minutes at 4°C and placed immediately on ice. The polar (methanol/water) layer and non-polar (chloroform) layers were subsequently transferred to 1.5 mL protein low binding microcentrifuge tubes. These metabolite suspensions were lyophilized overnight without heat in a ThermoScientific™ Savant™ ISS110 SpeedVac™ (Waltham, MA, USA) and stored at -80°C until the samples were shipped to Metabolon for further analysis. The remaining interphase layer was flash-frozen in liquid nitrogen and stored at -80°C until RNA extraction.

##### Metabolite Detection, Identification, and Quantification

All samples were analyzed by Metabolon (Morrisville, NC, USA) using four ultra-high-performance liquid chromatography/tandem accurate mass spectrometry (UHPLC/MS/MS) methods. The following is a summary of Metabolon’s procedure. All methods utilized a Waters ACQUITY ultra-performance liquid chromatography (UPLC) and a Thermo Scientific Q-Exactive high resolution/accurate mass spectrometer interfaced with a heated electrospray ionization (HESI-II) source and Orbitrap mass analyzer operated at 35,000 mass resolution. Lyophilized samples were reconstituted in compatible solvents containing a series of standards at fixed concentrations to assay consistency. Three aliquots were gradient eluted from a C18 column (Waters UPLC BEH C18-2.1x100 mm, 1.7 µm) using water and methanol, containing 0.05% perfluoropentanoic acid (PFPA) and 0.1% formic acid to detect hydrophilic compounds, methanol, acetonitrile, water, 0.05% PFPA and 0.01% formic acid for hydrophobic compounds, and methanol and water with 6.5 mM ammonium bicarbonate at pH 8 to for basic compounds. A fourth aliquot was analyzed via negative ionization following elution from a HILIC column (Waters UPLC BEH Amide 2.1x150 mm, 1.7 µm) using a gradient consisting of water and acetonitrile with 10mM ammonium formate, pH 10.8. The MS analysis alternated between MS and data-dependent MS scans using dynamic exclusion. The scan range varied slightly between methods but covered 70-1000 m/z. Controls included a pooled matrix sample that served as a technical replicate throughout the data set, extracted water samples as process blanks; and a cocktail of quality control (QC) standards.

Raw data was extracted, peak-identified and QC processed using Metabolon’s hardware and software. Compounds were identified by comparison to library entries of purified standards or recurrent unknown entities. Biochemical identifications are based on three criteria: retention index within a narrow RI window of the proposed identification, accurate mass match to the library +/- 10 ppm, and the MS/MS forward and reverse scores between the experimental data and authentic standards. The resulting data was then normalized to sample weight to account for differences in metabolite levels due to differences in the amount of material present in each sample. Finally, near zero variance metabolites were excluded from the dataset, and all metabolite features were centered and scaled to unit variances (80).

#### Microbiome profiling of wound-colonizing bacteria

##### 16S rRNA and cDNA isolation

16S rRNA was purified from the wound debridement samples utilizing the Qiagen RNeasy PowerBiofilm Kit (Germantown, MD, USA). The interphase layer from the metabolite extraction was resuspended in the supplied lysis buffer supplemented with 1% (v/v) of β-mercaptoethanol and homogenized, as previously described. Following lysis, the RNA was extracted and purified according to the manufacturer’s instructions. The integrity of the extracted RNA was evaluated on an Agilent 2200 Tape Station (Santa Clara, CA, USA). cDNA was generated from the extracted 16s rRNA using the Applied Biosystems High-Capacity cDNA Reverse Transcription Kit with RNAse Inhibitor according to manufacturer’s instructions.

##### 16S rRNA sequencing

The V1-V3 region of bacterial 16S rRNA gene was amplified from extracted tissue using Bac_8F and Bac_529r primers, Taq polymerase (Takara Biotechnology), and standard PCR cycling: 94^0^C for 30 secs followed by 30 cycles denaturation at 94^0^C for 30 secs, annealing at 52^0^C for 30 sec, and extension at 72^0^C for 30 seconds; followed by a final extension at 72^0^C for 10 mins with adaptor sequences for Illumina sequencing. The final library was pair-end sequenced (2 x 300 bp) on an Illumina MiSeq at the Center for Biofilm Engineering at Montana State University, using the MiSeq reagent kit v3, 600 cycles.

##### Taxonomic and abundance analysis of sequence data

The resulting sequence data were analyzed using QIIME2-2022.11 (81), running in Ubuntu 22.04.1. The DADA2 package (82) wrapped inside the QIIME2 pipeline was used for denoising and constructing amplicon sequence variants (ASVs). Reverse reads were truncated at 257 bp to remove low quality (q<30) base calls. Forward and reverse reads were trimmed by 20 and 15 bp, respectively, to remove primer sequences. Samples with a high percentage of chimeric sequences, low-quality reads, and those that had fewer than 300 hits were removed. Taxonomic analysis was performed using a pre-trained Bayesian naive classifier (gg-13-8-99-nb-classifier), which was trained on the Greengenes 16S rRNA gene database (83). Classified bacterial taxa down to genus level were used for input into the statistical model. Taxa with near zero variance, or whose counts amounted to less than 1% of the total were excluded from analysis. All remaining microbiome features underwent the centered log transformation (84) and were then standardized to zero means and unit variances (80).

### Integrative Model

#### Data

Following clinical annotation of the collected samples and data collection, three distinct datasets were compiled and herein referred to as the metabolome (metabolomics dataset), microbiome (16S rRNA sequencing data), and clinical markers (patient metadata). Samples within the compiled dataset were classified according to whether they came from healing versus non-healing wounds.

### DIABLO modeling

The healing outcome was modeled as a function of metabolome, microbiome, and clinical datasets using DIABLO (84–86), parameterized with between-dataset weights of 0.1 and two components per dataset. The weight values prioritize finding a model fit that discriminates between outcome classes but still learns inter-dataset correlations (84).

The number of features to select into the model for each dataset and component was determined by testing all possible combinations of feature numbers, subject to two constraints. First, single dataset modeling of the outcome using sPLS-DA established better predictive performance when a minimum of 80 and two features were selected from the metabolome and microbiome datasets, respectively (see Supplemental Table 2). Second, the maximum number of features that could be selected for each dataset was always half the number of features available after preprocessing. This restriction enforced significant regularization and acted as a hedge against overfitting. In all, 6,120 candidate DIABLO models were fit, representing all possible combinations of selected numbers of features in each dataset, on each component. Seven-fold cross-validation with 100 repeats was used to identify the final model — i.e., the fit with the lowest predictive error. After the DIABLO model was fit, the selected metabolite and microbiome features underwent a series of analyses to determine data set spefic trend.

### Feature Characterization

For the metabolomic set enrichment analysis (MSEA), the selected metabolite features were segregated based on expression level in the healing and non-healing groups. The segregated data sets were then analyzed independently in the Enrichment Analysis and Pathway Analysis module in MetaboAnalyst 6.0 (Ste. Anne de Bellevue, Quebec) (88). For the MSEA, overrepresentation analysis (ORA) was used to test if the metabolite set was represented more than expected by chance when compared to the Small Molecule Pathway Database (SMPDB) (88) with Bonferroni corrected P-values of p < 0.5 denoting significance. The 180 taxa fed into the DIABLO model were used to create a phylogenetic heat trees with differential abundances shown with heat map colors using the R package Metacoder (89).

### Statistics

R version 4.5.1 was used for statistical analysis (90). Continuous data are expressed as mean ± SEM. Categorical data are expressed as N (%). Continuous data comparisons in the clinical marker data set were performed using either a Welch’s t-test or a using the Wilcoxon rank sum test depending on data normality. The Fisher’s exact test was performed for categorical data. For metabolomics and microbiome data sets a mixed effects model with t-tests using Satterthwaite’s approximation. P-values were Benjamini-Hochberg adjusted with p < 0.05 considered significant. In the MSEA, ORA using a Bonferroni corrected P-values of p < 0.5 was used to denote significance.

### Data Availability

Global metabolomics data, 16s microbiome data, deidentified clinical metadata, and data processing code will be made available upon peer review publication at https://github.com/ammonslab/wound_healing_omics.

### Author contributions

Conceptualization, study design and supervision were done by CBA and MCBA. MMD supervised patient consent and sample collection. CBA, HLS, and TMWL processed patient samples. CH sequenced 16S rRNA samples. Data collation and preliminary data analysis was performed by CBA, CH, and MCD. JB designed and implemented the integrative model. The manuscript was written by CBA, JB, and MCBA, with edits from all other authors.

### Funding support

This work was supported in part by the U.S. Department of Veterans Affairs, Office of Research and Development Biomedical Laboratory Research Program Merit Award I01BX005244 (PI Ammons), a Veterans Affairs contract for sequencing and data analysis (VA SeqCURE – LSI Ammons), NIH grant 1KO1GM103821-01 (PI Ammons), NIH grant R03AI135998 (PI Ammons), the Idaho INBRE Program (NIH NIGMS P20 GM103408 - PD Bohach), and Project Four (PI Ammons) of the Idaho Center of Biomedical Research Excellence in Emerging/Reemerging Infectious Diseases (NIH NIGMS P20GM109007 – PD Stevens).

## Acknowlegements

None

