## Supplemental Table_Figures for "Integration of localized microbiome, metabolome, and clinical datasets predicts healing in chronic wounds among veterans"

### Supplemental material

Supplemental Table 1: Number of features selected in the final DIABLO model by dataset and component.

| Dataset | Measured | Modeled | Component one Selected | Component two Selected | Unique Selected |
| --- | --- | --- | --- | --- | --- |
| Metabolome | 865 | 854 | 80 | 80 | 153 |
| Microbiome | 634 | 180 | 2 | 6 | 8 |
| Clinical | 21 | 21 | 10 | 10 | 15 |
| Total | 1,520 | 1,055 | 92 | 96 | 176 |

Supplemental Table 2: Number of features selected and cross-validated errors for the best-performing single dataset sPLS-DA models.

|  | N Features | |  |
| --- | --- | --- | --- |
| Model | Component one | Component two | Error |
| Metabolome | 80 | 80 | 0.24 |
| Microbiome | 2 | 20 | 0.11 |
| Clinical | 10 | 10 | 0.02 |

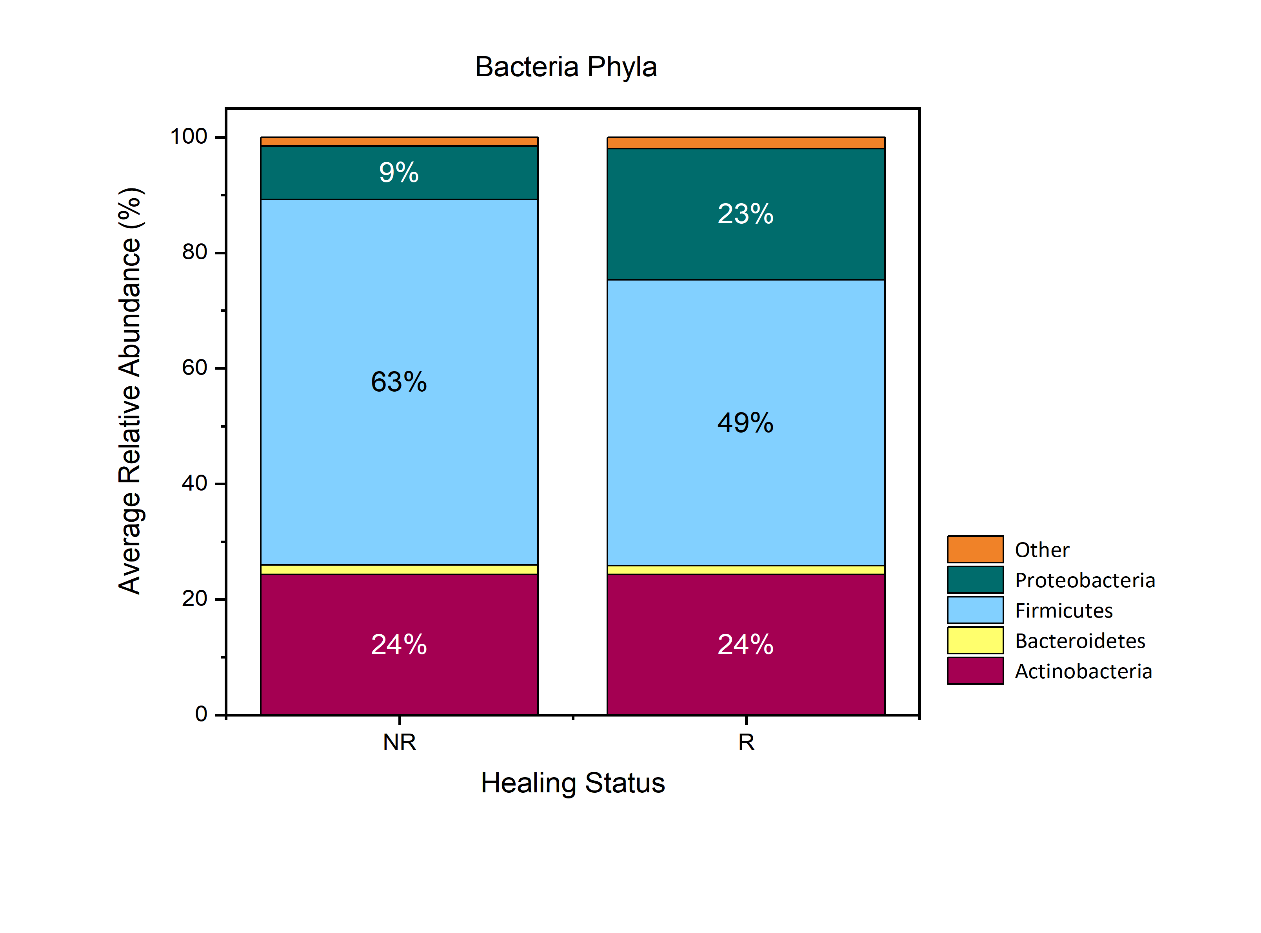

Non-Healers

Healers

Healing Status

Supplemental Figure 1: Stacked bar-chart showing relative abundance of major Phyla in Healers versus Non-Healers. Any phyla with mean abundances less than 1.5% were grouped into “other” category.
